# Prestimulus alpha-band power biases visual discrimination confidence, but not accuracy

**DOI:** 10.1101/089425

**Authors:** Jason Samaha, Luca Iemi, Bradley R. Postle

## Abstract

Oscillations in the alpha-band (8-13 Hz) of human electroencephalographic (EEG) recordings are thought to reflect cortical excitability. As such, the magnitude of alpha power prior to the onset of a near threshold visual stimulus has been shown to predict stimulus detectability. Mechanistically, however, non-specific increases in visual-cortical excitability should result in amplified signal as well as amplified noise, leaving actual discriminability unchanged. Using a two-choice orientation discrimination task with equally probable stimuli, we found that discrimination accuracy was unaffected by fluctuations in prestimulus alpha-band power. Decision confidence, on the other hand, was strongly negatively correlated with prestimulus alpha power. This finding constitutes a clear dissociation between objective and subjective measures of visual perception as a function of prestimulus cortical excitability. This dissociation is predicted by models of perceptual confidence under which the balance of evidence in favor of each choice drives objective performance but only the magnitude of evidence in favor of the chosen stimulus drives subjective reports, suggesting that human perceptual confidence can be suboptimal.

## 1. Introduction

Excitability of the visual cortex has been directly linked to fluctuations in the power of alpha-band (8-13 Hz) oscillations in human electroencephalographic (EEG) recording (Brandt & Jansen, 1991; Rajagovindan & Ding, 2010; Romei, Brodbeck, et al., 2008; Romei, Rihs, Brodbeck, & Thut, 2008; Samaha, Gosseries, & Postle, 2016). Accordingly, recent work has found that variability in the detection of near-threshold visual stimuli is explained by variability in alpha-band power just prior to stimulus onset. A now-typical finding is that the probability of detecting a visual stimulus increases when prestimulus alpha power is low (Babiloni, Vecchio, Bultrini, Romani, & Rossini, 2006; Busch, Dubois, & VanRullen, 2009; Dijk, Schoffelen, Oostenveld, & Jensen, 2008; Ergenoglu et al., 2004; Mathewson, Gratton, Fabiani, Beck, & Ro, 2009; Romei, Gross, & Thut, 2010). From a Signal Detection Theory (SDT) framework, however, an increased probability of detection (i.e., hit rate) could result from change either in perceptual sensitivity (i.e., *d’*) or in response criterion (Green & Swets, 1966). That is, when alpha power is low, observers may be better able to discriminate a visual stimulus from noise or they may be more likely to report seeing a stimulus, regardless of whether one was actually present.

A recent experiment confirms the latter scenario. Limbach & Corballis (2016) performed a SDT analysis of detection performance as a function of prestimulus alpha levels. They observed that response criterion, and not *d’,* was related to prestimulus alpha, such that observers adopted a more conservative criterion when alpha power was high. However, this finding leaves open the question of how prestimulus alpha changes subjective and objective responses in a discrimination task with equally probable stimuli, where criterion is presumably balanced (Macmillan & Creelman, 2004). Because changes in criterion have been linked to changes in subjective awareness reports (Peters, Ro, & Lau, 2016; Rahnev et al., 2011), we hypothesize that prestimulus alpha may impact confidence ratings in a discrimination task, but should not affect discrimination accuracy. This hypothesis is motivated by the idea that when cortical excitability is non-specifically increased, neurons representing the presented stimulus (correct choice) as well as those representing the non-presented alternative (incorrect choice) should both increase their firing rates by the same amount, leaving discriminability between the two unaffected. However, if confidence is driven primarily by evidence in favor of the choice stimulus, rather than the balance of evidence for both stimuli (e.g., Maniscalco, Peters, & Lau, 2016), then confidence will be systematically higher when cortical excitability is higher (e.g., when alpha is low), despite no change in accuracy. This hypothesis is depicted within a SDT framework in Fig. 1.

**Fig. 1.**
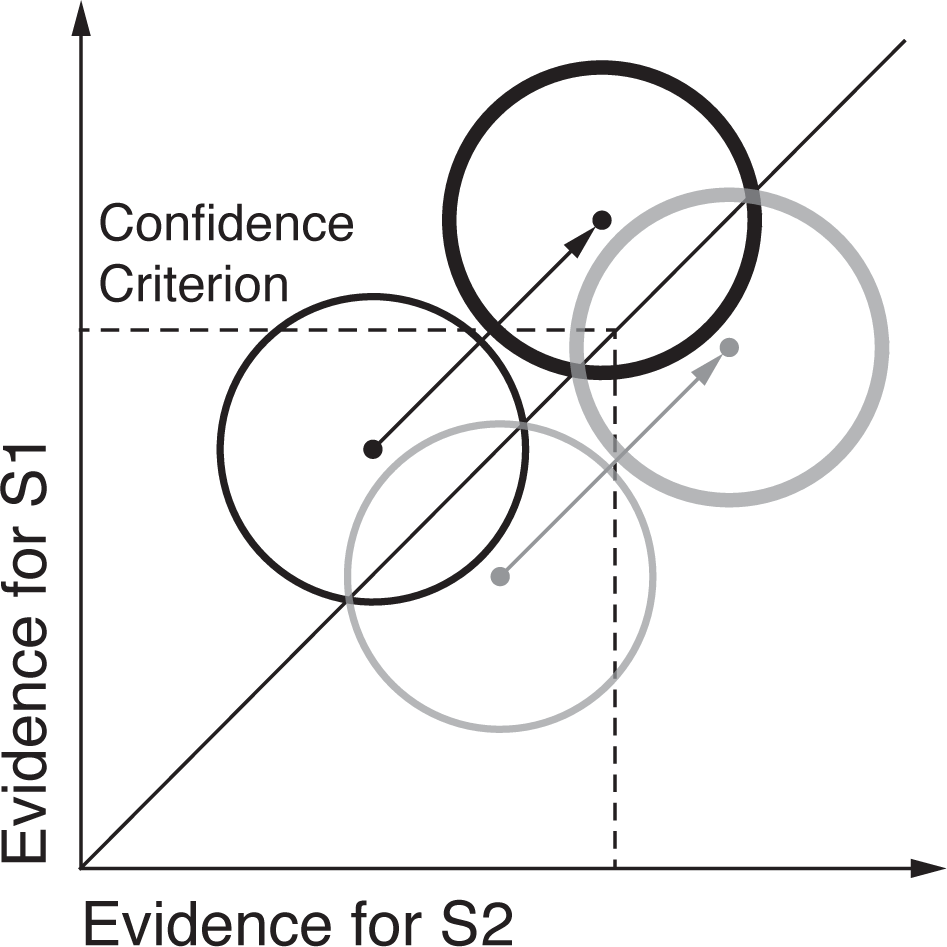
In the two-dimensional SDT model for discrimination, a decision is made regarding which of two stimuli was presented (S1 or S2; black and grey lines, respectively) using evidence (e.g., the firing rates of neurons representing S1 and S2) sampled from bivariate Gaussian distributions. The diagonal line represents the optimal decision criterion: if evidence on a given trial falls to the left of the diagonal, S1 is chosen, otherwise, S2. Thus, discrimination accuracy is determined by the separation between the black and grey distributions with respect to the diagonal. Rating confidence according to the amount of evidence for the chosen stimulus, rather than the balance of evidence for both, results in confidence criteria (dashed lines) marked at various points along the x and y axes. We hypothesize that during states of high cortical excitability both evidence distributions are translated diagonally (thick lines), reflecting increased evidence for both S1 and S2. The dissociation between confidence and accuracy comes about because the two pairs of distributions are identical with respect to the decision boundary (diagonal), but not with respect to the confidence criterion. This predicts higher confidence when visual cortical excitability is high (i.e., alpha is low), but no change in accuracy. For a more detailed treatment and empirical evidence in support of this model, see Maniscalco, Peters, & Lau (2016). Note that the choice of a single confidence criterion and its location is arbitrary and used here only for illustration.

Recent psychophysical studies have borne out the proposal that confidence is disproportionality affected by evidence in favor of a decision, rather than the balance of evidence between alternatives. For example, Zylberberg, Barttfeld, & Sigman (2012) continuously varied the luminance of two stimuli as observers decided which was brightest. Their findings show that, whereas choice accuracy was determined by relative difference in luminance between the two stimuli, confidence was insensitive to fluctuations in luminance for the non-chosen stimulus but was driven by the absolute luminance of the chosen stimulus. Subsequent work found that proportionally increasing the contrast of a target as well as the contrast of noise (or a non-target, e.g., Koizumi, Maniscalco, & Lau, 2015; Expt 1A) led to increased confidence despite no change to accuracy (Koizumi et al., 2015; Samaha, Barrett, Sheldon, LaRocque, & Postle, 2016). Here, we measured prestimulus alpha power as a trial-by-trial index of cortical excitability while observers judged the orientation of a grating and provided subjective confidence ratings. We found a robust negative relationship between prestimulus alpha power and confidence ratings, but no relationship to accuracy, suggesting that states of high visual cortical excitability are associated with an enhanced sense of subjective confidence despite no change in actual performance. We view this finding as providing support for models of subjective awareness according to which the absolute value of evidence in support of a decision underlies confidence (Maniscalco et al., 2016; Paz, Insabato, Zylberberg, Deco, & Sigman, 2016; Zylberberg et al., 2012; Zylberberg, Roelfsema, & Sigman, 2014).

## 2. Materials and methods

### 2.1 Subjects

10 participants (5 female; age range 21-34) were recruited from the University of Wisconsin-Madison community and participated for monetary compensation. The UW-Madison Health Sciences Institutional Review Board approved the study protocol. All subjects provided informed consent and self-reported normal or corrected-to-normal vision and no history of neurological disorders.

### 2.2 Stimuli

Target stimuli were sinusoidal luminance gratings embedded in random dot noise presented within a circular aperture (see Fig. 2A). Gratings subtended 2.5 degrees of visual angle (DVA), had a spatial frequency of 1.5 cycles/DVA, a phase of zero, and were either rotated 45 or -45 degrees from vertical (randomly determined). Noise consisted of random black and white pixels. The contrast of the grating was determined for each subject by an adaptive staircase procedure (see *Procedure*). On a random half of the trials the contrast of both the signal and the noise was halved, which was not expected to impact accuracy (Samaha, Barrett, et al., 2016), but this manipulation was not explored here. Stimuli were presented on an LCD screen (34 cm; 2560 x 1600; 60 Hz refresh rate) that was viewed at a distance of 63 cm in a dimly lit room. Stimuli were generated and presented using the MGL toolbox (http://gru.stanford.edu) running in MATLAB 2015b (MathWorks, Natick, MA). Fixation (a light grey point, 0.08 DVA) was centered on the screen and all stimuli were presented centrally against a grey screen background.

**Fig. 2.**
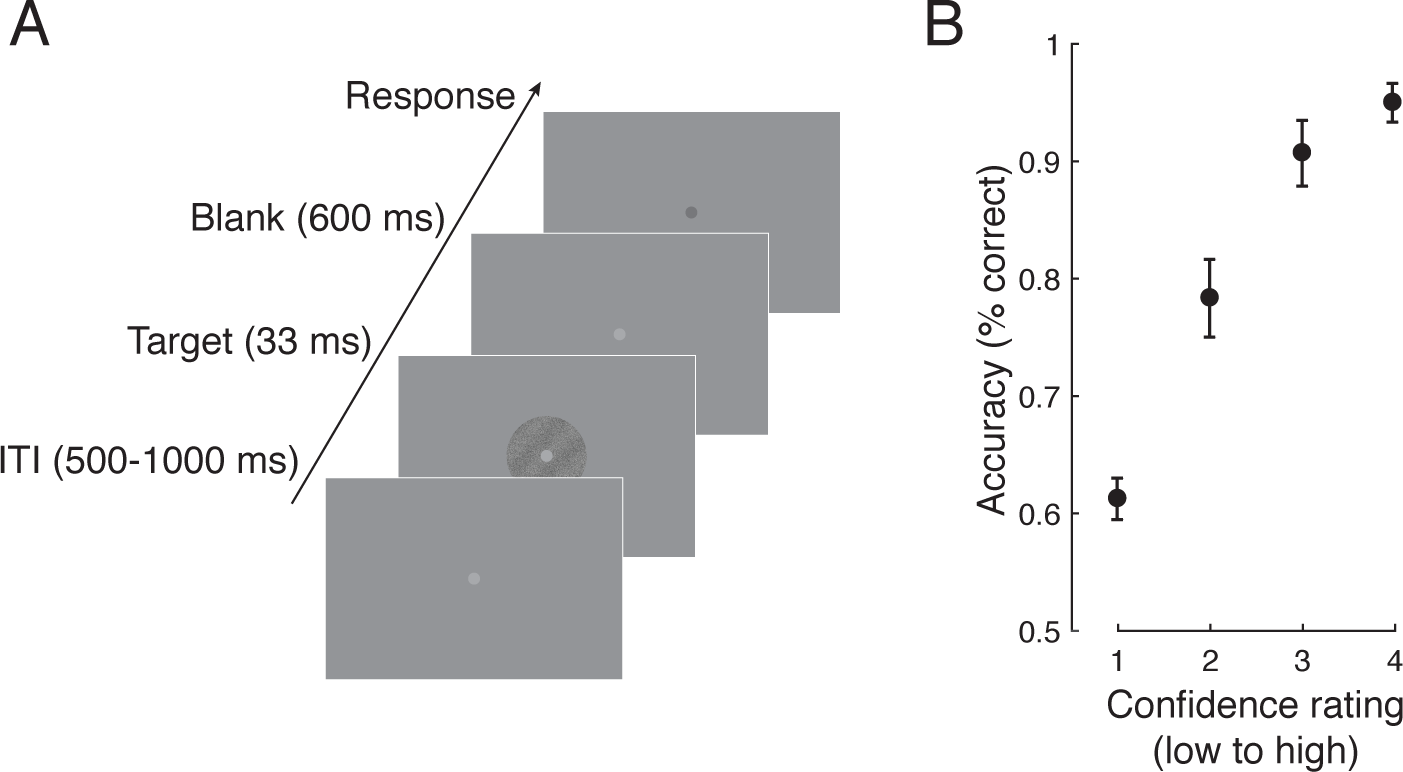
Task design and performance. **A.** On each trial a threshold-level grating embedded in noise was centrally presented and subjects indicated its tilt (-45^o^ or 45^o^ from vertical, equally likely) followed by a rating of how confident they were in their judgment (1-4) while EEG was recorded. **B.** Discrimination accuracy showed a positive monotonic relationship with subjective confidence (N=10,695 trials), indicating that the confidence scale was used appropriately. Analysis of the EEG focused on the inter-trial interval (ITI) just prior to target onset. Error bars indicate ± SEM.

### 2.3 Procedure

Each subject completed 1,230 trials of a two-choice orientation discrimination task (Fig. 2A) in which they first decided whether the target grating was rotated left or right of vertical and then provided a confidence rating (1-4; from guessing to highly confident). An adaptive 1-up 3-down staircase procedure controlled by the PEST algorithm (Taylor & Creelman, 1967) was run for the first 110 trials. This was intended to produce ~ 79% discrimination accuracy. Each trial (Fig. 2) began with a central fixation point presented for a random duration between 500 and 1000 ms, followed by a grating for 33 ms, followed by fixation again, which dimmed slightly after 600 ms to indicate that responses could be made with no time limit. This 600 ms grace period was included to allow the evoked response in the EEG to unfold without motor contamination and to ensure that subjects had accumulated the same amount of evidence for both the orientation and confidence judgment (Navajas, Bahrami, & Latham, 2016). Note that this precluded the study of reaction times. Once fixation dimmed, subjects indicated their choice using their right hand on the left and right arrows of a computer keyboard followed by their confidence using their left hand on the number keys 1-4. The remaining 1,120 task trials were procedurally identical except that the contrast of the grating was fixed to be the average of the last 6 reversals found during the staircase. These trials were split into 7 blocks with short rests in between. Total experiment time, including EEG setup, was approximately 2.5 hrs.

### 2.4 EEG acquisition and preprocessing

EEG was collected from 60 Ag/AgCl electrodes (Nexstim, Helsinki, Finland), each with impedance kept below 10 kΩ. A single electrode placed on the forehead was used as the reference, and eye movements were recorded with two biploar electrodes placed to the side, and underneath, of the right eye. Data were acquired at a rate of 1,450 Hz with 16-bit resolution. EEG was processed offline with custom MATLAB scripts (version R2014b) and with the EEGLAB toolbox version 13.5 (Delorme & Makeig, 2004). Recordings were visually inspected and noisy channels (1.4 on average) were spherically interpolated. Data were downsampled to 250 Hz and re-referenced to the median of all electrodes. A one-pass zero-phase Hamming windowed-sinc FIR filter between 0.5 and 50 Hz was applied to the data (EEGLAB function *pop_eegfiltnew.m*) and epochs spanning -1300 to 1300 relative to grating onset were extracted (time-stamped with a photodiode recording made during stimulus presentation). A prestimulus baseline of -200 to 0 ms was subtracted from each trial. Individual trials were visually inspected and those containing excessive noise, muscle artifacts, or ocular artifacts occurring contemporaneously with target onset were removed (30 - 82 trials per subject). These trials were also excluded from the analysis of behavioral data. Independent components analysis using the INFOMAX algorithm (EEGLAB function *binica.m*) was used to subtract remaining ocular artifacts not coinciding with the stimulus (2.8 components on average).

### 2.5 Time-frequency analysis

To estimate power across time and frequencies, data from each channel and trial were convolved with a family of complex Morlet wavelets spanning 2–50 Hz in 1-Hz steps with wavelet cycles increasing linearly between three and eight cycles as a function of frequency. Power was obtained by squaring the absolute value of the resulting complex time series and an analysis epoch between -500 and 200 ms relative to stimulus onset was extracted. We focused our analysis on a cluster of posterior electrodes defined by visually selecting contiguous electrodes that showed the largest alpha power between -500 and 0 ms prior to stimulus onset when averaged across all trials, as displayed in Fig. 3A.

**Fig. 3.**
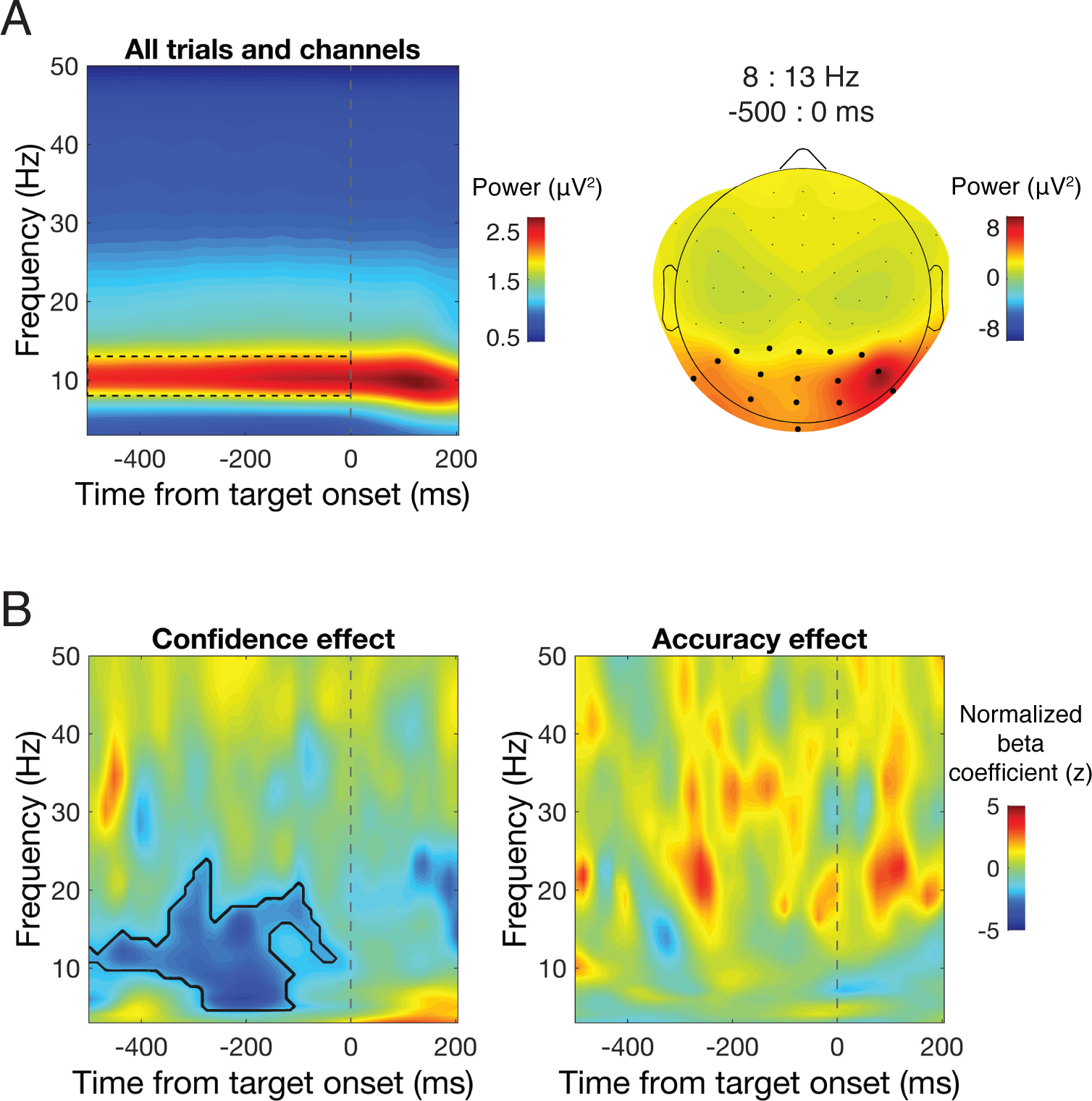
Single-trial regression of time-frequency power. **A.** Time-frequency power averaged over all trials and electrodes revealed a clear peak in alpha-band power during the prestimulus window (left; grey dashed box), which was maximal over posterior electrodes (right). Black dots denote the electrode cluster used for subsequent analysis. **B.** The results of a single-trial regression of confidence (left) and accuracy (right) on power across a range of times and frequencies revealed that prestimulus low-frequency power prior to target onset was negatively correlated with confidence ratings, but not discrimination accuracy. Black contour denotes significant cluster-corrected effects at p<0.05.

### 2.6 Single-trial regression

To relate single-trial estimates of power across time and frequency space to single trial behavioral responses (accuracy and confidence), we used a non-parametric single-trial multiple regression approach. Whereas traditional analyses in cognitive electrophysiology proceed by averaging over trials in a condition in order to reduce noise associated with single-trial data, this approach does not account for trial-by-trial variance. Because we collected a large number of trials in each subject, we incorporated this single-trial information in a multilevel model. For each time-point, frequency, channel (in our region of interest), and subject, regression coefficients that describe the relationship between power and accuracy and between power and confidence were estimated according to the linear model:

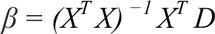

where *X* is the design matrix containing one column for the intercept (all ones), one column of accuracy scores (dummy coded as 1 or 0), and one column of confidence ratings across all trials. *^T^* and ^−1^ indicate the matrix transpose and inverse, and *D* is the vector of power data from all trials at a given time-frequency point and channel. Power values were first rank scored to mitigate the influence of outliers (this is equivalent to a Spearman correlation), and *β* was estimated using a least-squares solution. The resulting beta coefficients describe the independent contributions of accuracy and confidence to explaining prestimulus power. Beta coefficients were then converted into a *z*-statistic relative to a subject-specific null hypothesis distribution obtained by repeatedly shuffling the mapping between the data and the design matrix (see *Statistics*). The resulting *z*-statistics were averaged over all electrodes in our region of interest.

### 2.7 Alpha power binning analysis

We additionally binned confidence and accuracy into 10 deciles according to prestimulus alpha power levels obtained from a discrete fast Fourier transform (FFT) of prestimulus data. This analysis was motivated by three considerations. 1) Because time-frequency analysis necessarily results is some temporal smoothing, it is possible that a poststimulus effect can smear in time and appear in the prestimulus window (Zoefel & Heil, 2013). We therefore extracted epochs between -500 and 0 ms from each trial and estimated power with a zero-padded (resolution: 0.33 Hz), Hamming window-tapered, FFT, which ensured that only prestimulus data was included in the analysis. 2) Substantial variability in peak alpha frequency exists across individuals (Haegens, Cousijn, Wallis, Harrison, & Nobre, 2014) that was not taken into account in the time-frequency analysis. For this reason, we binned confidence and accuracy according to alpha power defined as the average over a ± 2 Hz window centered on each individuals alpha peak (identified from the power spectrum averaged across channels in our posterior region of interest). 3) Lastly, this binning analysis served as a more traditional approach by which to verify the single-trial regression results.

### 2.8 Statistics

Level one (subject-level) statistics were performed for the singe-trial time-frequency regression analysis by randomly permuting the mapping between confidence/accuracy and neural data 2000 times and recomputing beta coefficients each time. The coefficient associated with the true data mapping was then converted to a *z*-statistic relative to the mean and standard deviation of the permuted data, which was then averaged over posterior channels. This resulted in a *z* value for each subject and time-frequency point. This approach incorporates knowledge about variability in the subject-level effects into the subsequent group-level analyses. Level two (group-level) statistics and significance values were computed by means of non-parametric permutation tests in combination with cluster correction to address multiple comparisons across time and frequency points. To estimate group-level null hypothesis distributions, on each of 5000 permutations, z-scores from a random subset of subjects for both the confidence and accuracy effect were multiplied by -1 and a two-tailed *t*-test against zero was computed (this is equivalent to randomly swapping the order of the condition subtraction, e.g., A-B vs. B-A, (Maris & Oostenveld, 2007). On each permutation, the size of the largest contiguous cluster of significant (α = 0.05) time-frequency points was saved, forming a distribution of cluster sizes expected under the null hypothesis. The *t*-statistics associated with the true group-level test were converted to *z*-statistics relative to the mean and standard deviation of the empirical null hypothesis distributions and only clusters of significant *z* values (*z*_crit_ = 1.96) exceeding the 95% percentile of the distribution of cluster sizes expected under the null hypothesis were considered significant.

Regarding the statistical analysis of the binned data, we computed correlation coefficients at the group level between alpha power bin and accuracy/confidence. Additionally, we fit linear regression slopes to each subject’s accuracy by alpha and confidence by alpha distributions and compared these slopes against zero at the group level with a two-tailed *t*-test.

## 3. Results

### 3.1 Behavioral performance

On average, performance on the discrimination task with 78.8% accuracy (SEM = 2.8%), indicating that the staircase procedure was effective. Accuracy increased monotonically as a function of confidence (Fig. 2B), and confidence was significantly higher on correct trials (*M* = 2.35, SEM = 0.18) compared to incorrect trials (*M* = 1.64, SEM = 0.14; t(1,9) = 4.57, p = 0.001), suggesting that the confidence scale was used appropriately. All subjects showed this pattern (not shown).

### 3.2 Prestimulus alpha power predicts confidence but not accuracy

Prestimulus alpha power was largest over a cluster of occipital-parietal electrodes (Fig. 3A), which we aggregated over for further analysis. We found a significant negative relationship between confidence and prestimulus power between 5 and 20 Hz, from -500 ms until stimulus onset, indicating that low prestimulus power in this time-frequency range was associated higher subsequent confidence ratings (Fig. 3B, left). This effect was centered on the traditional range of alpha (8-13 Hz) but notably included frequencies just above and below this range. In contrast, no significant effect of power on accuracy was found in any of the time-frequency ranges examined here (Fig. 3B, right). Binning accuracy and confidence according to prestimulus alpha power revealed a strong negative correlation between alpha and confidence ratings (r = -0.84, p = 0.002; Fig. 4), indicating that prestimulus alpha power accounts for 71% of the variance in confidence ratings at the group level. The correlation between prestimulus alpha and accuracy was non-significant (r = -0.39 p = 0.26). Supporting this pattern, slopes fit to individual subject’s data relating alpha power to confidence were significantly different from zero (mean slope = -0.012, t(1,9) = -3.12, p = 0.012), but slopes relating alpha to accuracy were not (mean slope = -0.002, t(1,9) = -1.38, p = 0.201). Accuracy-power and confidence-power slopes were also significantly different from each other (t(1,9) = 2.80, p = 0.021), indicating that prestimulus alpha power predicts confidence to a greater extent than it predicts accuracy.

**Fig. 4.**
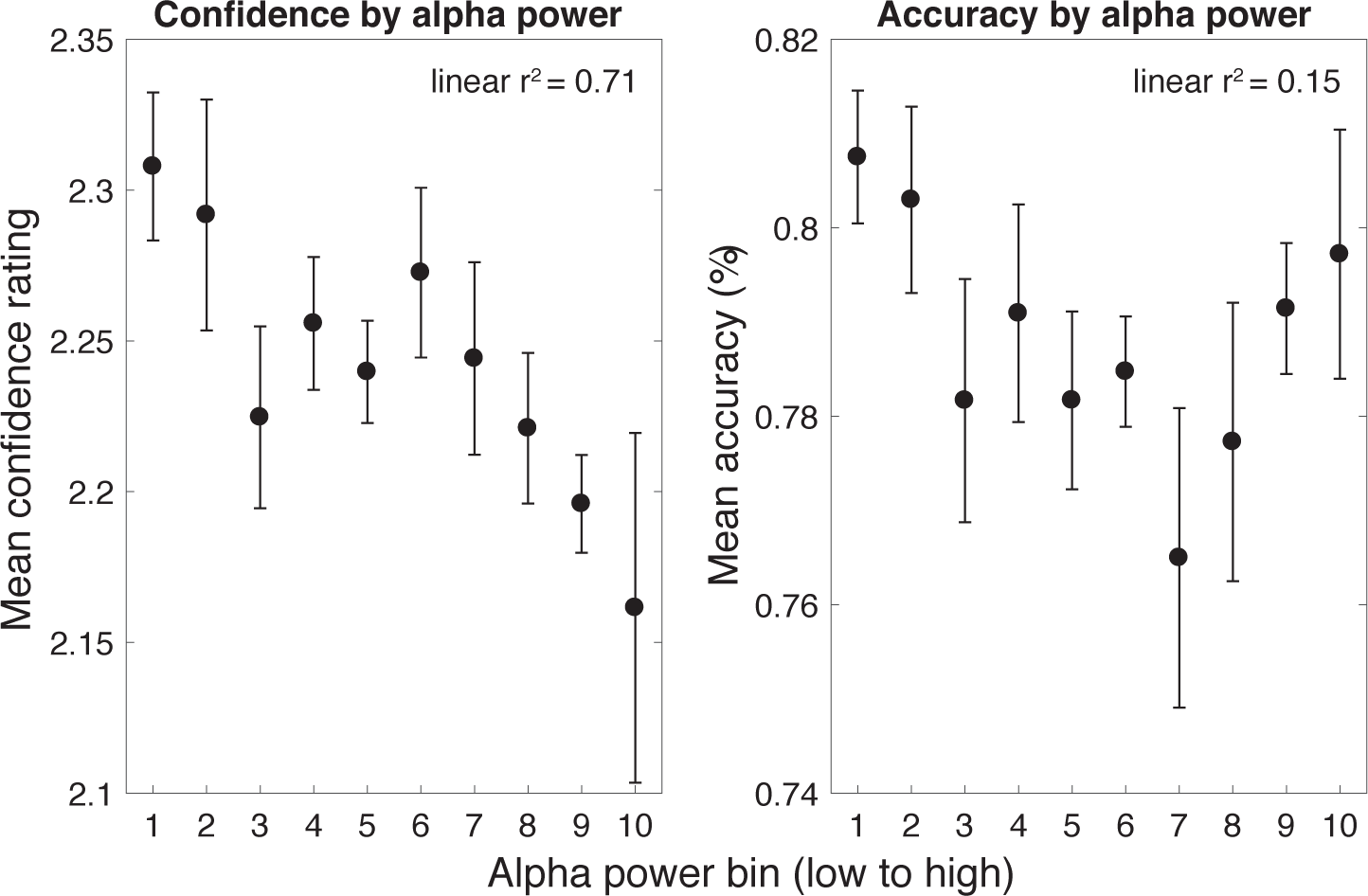
Confidence and accuracy as a function of prestimulus alpha power. FFT analysis of the prestimulus window (-500 : 0 ms) using individual peak alpha frequencies revealed the same pattern of effects: prestimulus alpha power was strongly negatively correlated with confidence but not accuracy. Error bars indicate ± within-subject SEM (Morey, 2008).

## 4. Discussion

Using a two-choice orientation discrimination task with equally probable stimuli, we found that fluctuations in prestimulus alpha-band power were independent of discrimination accuracy, but were strongly negatively related to subjective confidence. This was revealed from a single-trial regression analysis jointly accounting for the effects of accuracy and confidence on time-frequency power (Fig. 3B) as well as when confidence and accuracy were binned according to prestimulus power at each individual’s peak alpha frequency (Fig. 4).

Alpha-band oscillations have been tightly linked to cortical excitability in humans (Mathewson et al., 2009; Rajagovindan & Ding, 2010; Romei, Brodbeck, et al., 2008; Romei et al., 2010; Romei, Rihs, et al., 2008; Samaha, Gosseries, et al., 2016), in non-human primates (Haegens, Nácher, Luna, Romo, & Jensen, 2011), and in biophysical neural models (Becker, Knock, Ritter, & Jirsa, 2015; Lundqvist, Herman, & Lansner, 2013). We suggest that non-specific spontaneous fluctuations in visual cortical excitability should manifest as fluctuations in neural responses representing the presented (correct) stimulus as well non-presented (incorrect) alternative, which should result in no change to visual discrimination ability. Subjective confidence, on the other hand, may only be sensitive to the absolute amount of evidence in favor of a perceptual choice (Maniscalco et al., 2016; Zylberberg et al., 2012), which would result in higher confidence when the excitability is higher (Fig. 1). Whereas this has been empirically demonstrated through manipulations of stimulus evidence (Koizumi et al., 2015; Samaha, Barrett, et al., 2016; Zylberberg et al., 2014), we show that this is the case when internal evidence fluctuates spontaneously as a function of cortical excitability, as indexed by prestimulus alpha power. Although numerous studies have observed a relationship between prestimulus alpha power and hit rates on visual detection tasks (Busch et al., 2009; Dijk et al., 2008; Ergenoglu et al., 2004; Mathewson et al., 2009; Romei et al., 2010), it has recently been demonstrated that these findings could result from a shift in criterion, rather than perceptual sensitivity (Limbach & Corballis, 2016). In that study, when alpha power was low, observers more frequently reported stimulus presence. Our findings support this result in that we found that observers more readily rate higher confidence when prestimulus alpha is low, despite no change in task performance. Modulation of prestimulus alpha power has, however, been associated with increased perceptual sensitivity in the context of spatial cueing tasks where it is commonly observed that preparatory alpha increases over electrodes contralateral to the attended portion of visual space (Foxe & Snyder, 2011; Kelly, Lalor, Reilly, & Foxe, 2006; Samaha, Sprague, & Postle, 2016; Sauseng et al., 2005). In these paradigms, the spatially-specific decrease/increase of alpha power in task-relevant/-irrelevant areas may selectively boost the representation of relevant sensory information and attenuate the representation of distracting information, thereby inducing performance enhancements (Jensen & Mazaheri, 2010). In contrast, our results pertain to scenarios where cortical excitability undergoes a non-specific change, as reflected by spontaneous fluctuations of alpha power.

Our findings bare resemblance to a recent functional magnetic resonance imaging experiment where spontaneous fluctuations in hemodynamic activity in the dorsal attention network (DAN) prior to the onset of a dot motion stimulus was shown to relate to subjective confidence, but not objective discrimination (Rahnev, Bahdo, Lange, & Lau, 2012). As regions of the DAN have been implicated as substrates of attentional control underlying modulation of posterior alpha oscillations (Capotosto, Babiloni, Romani, & Corbetta, 2009; Marshall, O’Shea, Jensen, & Bergmann, 2015; Mathewson et al., 2014), it is possible that the relationship between alpha power and confidence we observed is itself related to fluctuations of activity in attentional control networks.

Importantly, our findings constitute a clear dissociation between objective and subjective measures of perception. The ability to induce such a dissociation has been argued to be an important tool for teasing apart the functional and neural basis of subjective awareness while controlling for basic task performance confounds (Lau & Passingham, 2006; Lau & Rosenthal, 2011; Samaha, 2015). Here, we show that it is possible to dissociate confidence from accuracy solely on the basis of prestimulus brain states measurable with EEG. This approach may compliment existing methods which rely on stimulus manipulations and may therefor also include low-level confounds (see e.g., Jannati & Di Lollo, 2012). Furthermore the existence of a dissociation between subjective and objective perception can help clarify models of subjective reports. In the SDT framework shown here, our results suggest that observers do not rate their confidence according to the balance of evidence for both choices in the way that the choice itself is made. Rather, confidence is primarily rated in accordance with the magnitude of evidence in favor of the chosen stimulus (Maniscalco et al., 2016; Paz et al., 2016; Zylberberg et al., 2012). Models whereby subjective confidence is an optimal statistical computation of the likelihood of having made a correct choice (e.g., Sanders, Hangya, & Kepecs, 2016), may have a hard time accommodating such a dissociation between decision accuracy and confidence.

## Acknowledgments

The authors would like to thank Dr. Bas Rokers and Dr. Niko Busch for very useful discussion.

## References

Babiloni, C., Vecchio, F., Bultrini, A., Romani, G. L., & Rossini, P. M. (2006). Pre- and Poststimulus Alpha Rhythms Are Related to Conscious Visual Perception: A High-Resolution EEG Study. Cerebral Cortex, 16(12), 1690–1700. https://doi.org/10.1093/cercor/bhj104

Becker, R., Knock, S., Ritter, P., & Jirsa, V. (2015). Relating Alpha Power and Phase to Population Firing and Hemodynamic Activity Using a Thalamo-cortical Neural Mass Model. PLOS Comput Biol, 11(9), e1004352. https://doi.org/10.1371/journal.pcbi.1004352

Brandt, M. E., & Jansen, B. H. (1991). The relationship between prestimulus-alpha amplitude and visual evoked potential amplitude. The International Journal of Neuroscience, 61(3–4), 261–268.

Busch, N. A., Dubois, J., & VanRullen, R. (2009). The Phase of Ongoing EEG Oscillations Predicts Visual Perception. The Journal of Neuroscience, 29(24), 7869–7876. https://doi.org/10.1523/JNEUROSCI.0113-09.2009

Capotosto, P., Babiloni, C., Romani, G. L., & Corbetta, M. (2009). Frontoparietal Cortex Controls Spatial Attention through Modulation of Anticipatory Alpha Rhythms. The Journal of Neuroscience, 29(18), 5863–5872. https://doi.org/10.1523/JNEUROSCI.0539-09.2009

Delorme, A., & Makeig, S. (2004). EEGLAB: an open source toolbox for analysis of single-trial EEG dynamics including independent component analysis. Journal of Neuroscience Methods, 134(1), 9–21. https://doi.org/10.1016/j.jneumeth.2003.10.009

Dijk, H. van, Schoffelen, J.-M., Oostenveld, R., & Jensen, O. (2008). Prestimulus Oscillatory Activity in the Alpha Band Predicts Visual Discrimination Ability. The Journal of Neuroscience, 28(8), 1816–1823. https://doi.org/10.1523/JNEUROSCI.1853-07.2008

Ergenoglu, T., Demiralp, T., Bayraktaroglu, Z., Ergen, M., Beydagi, H., & Uresin, Y. (2004). Alpha rhythm of the EEG modulates visual detection performance in humans. Cognitive Brain Research, 20(3), 376–383. https://doi.org/10.1016/j.cogbrainres.2004.03.009

Foxe, J. J., & Snyder, A. C. (2011). The role of alpha-band brain oscillations as a sensory suppression mechanism during selective attention. Perception Science, 2, 154. https://doi.org/10.3389/fpsyg.2011.00154

Green, D. M., & Swets, J. A. (1966). Signal Detection Theory and Psychophysics. Oxford, England: John Wiley.

Haegens, S., Cousijn, H., Wallis, G., Harrison, P. J., & Nobre, A. C. (2014). Inter- and intra-individual variability in alpha peak frequency. Neuroimage, 92(100), 46–55. https://doi.org/10.1016/j.neuroimage.2014.01.049

Haegens, S., Nácher, V., Luna, R., Romo, R., & Jensen, O. (2011). α-Oscillations in the monkey sensorimotor network influence discrimination performance by rhythmical inhibition of neuronal spiking. Proceedings of the National Academy of Sciences, 108(48), 19377–19382. https://doi.org/10.1073/pnas.1117190108

Jannati, A., & Di Lollo, V. (2012). Relative blindsight arises from a criterion confound in metacontrast masking: Implications for theories of consciousness. Consciousness and Cognition, 21(1), 307–314. https://doi.org/10.1016/j.concog.2011.10.003

Jensen, O., & Mazaheri, A. (2010). Shaping Functional Architecture by Oscillatory Alpha Activity: Gating by Inhibition. Frontiers in Human Neuroscience, 4. https://doi.org/10.3389/fnhum.2010.00186

Kelly, S. P., Lalor, E. C., Reilly, R. B., & Foxe, J. J. (2006). Increases in Alpha Oscillatory Power Reflect an Active Retinotopic Mechanism for Distracter Suppression During Sustained Visuospatial Attention. Journal of Neurophysiology, 95(6), 3844–3851. https://doi.org/10.1152/jn.01234.2005

Koizumi, A., Maniscalco, B., & Lau, H. (2015). Does perceptual confidence facilitate cognitive control? Attention, Perception & Psychophysics, 77(4), 1295–1306. https://doi.org/10.3758/s13414-015-0843-3

Lau, H. C., & Passingham, R. E. (2006). Relative blindsight in normal observers and the neural correlate of visual consciousness. Proceedings of the National Academy of Sciences, 103(49), 18763–18768. https://doi.org/10.1073/pnas.0607716103

Lau, H., & Rosenthal, D. (2011). Empirical support for higher-order theories of conscious awareness. Trends in Cognitive Sciences, 15(8), 365–373. https://doi.org/10.1016/j.tics.2011.05.009

Limbach, K., & Corballis, P. M. (2016). Prestimulus alpha power influences response criterion in a detection task. Psychophysiology. https://doi.org/10.1111/psyp.12666

Lundqvist, M., Herman, P., & Lansner, A. (2013). Effect of Prestimulus Alpha Power, Phase, and Synchronization on Stimulus Detection Rates in a Biophysical Attractor Network Model. The Journal of Neuroscience, 33(29), 11817–11824. https://doi.org/10.1523/JNEUROSCI.5155-12.2013

Macmillan, N. A., & Creelman, C. D. (2004). Detection Theory: A User’s Guide. Psychology Press.

Maniscalco, B., Peters, M. A. K., & Lau, H. (2016). Heuristic use of perceptual evidence leads to dissociation between performance and metacognitive sensitivity. Attention, Perception, & Psychophysics, 1–15. https://doi.org/10.3758/s13414-016-1059-x

Maris, E., & Oostenveld, R. (2007). Nonparametric statistical testing of EEG- and MEG-data. Journal of Neuroscience Methods, 164(1), 177–190. https://doi.org/10.1016/j.jneumeth.2007.03.024

Marshall, T. R., O’Shea, J., Jensen, O., & Bergmann, T. O. (2015). Frontal Eye Fields Control Attentional Modulation of Alpha and Gamma Oscillations in Contralateral Occipitoparietal Cortex. The Journal of Neuroscience, 35(4), 1638–1647. https://doi.org/10.1523/JNEUROSCI.3116-14.2015

Mathewson, K. E., Beck, D. M., Ro, T., Maclin, E. L., Low, K. A., Fabiani, M., & Gratton, G. (2014). Dynamics of Alpha Control: Preparatory Suppression of Posterior Alpha Oscillations by Frontal Modulators Revealed with Combined EEG and Event-related Optical Signal. Journal of Cognitive Neuroscience, 26(10), 2400–2415. https://doi.org/10.1162/jocn_a_00637

Mathewson, K. E., Gratton, G., Fabiani, M., Beck, D. M., & Ro, T. (2009). To See or Not to See: Pre-stimulus Alpha Phase Predicts Visual Awareness. The Journal of Neuroscience, 29(9), 2725–2732. https://doi.org/10.1523/JNEUROSCI.3963-08.2009

Morey, R. D. (2008). Confidence Intervals from Normalized Data: A correction to Cousineau (2005). Tutorials in Quantitative Methods for Psychology, 4, 61–64.

Navajas, J., Bahrami, B., & Latham, P. E. (2016). Post-decisional accounts of biases in confidence. Current Opinion in Behavioral Sciences, 11, 55–60. https://doi.org/10.1016/j.cobeha.2016.05.005

Paz, L., Insabato, A., Zylberberg, A., Deco, G., & Sigman, M. (2016). Confidence through consensus: a neural mechanism for uncertainty monitoring. Scientific Reports, 6. https://doi.org/10.1038/srep21830

Peters, M. A. K., Ro, T., & Lau, H. (2016). Who’s afraid of response bias? Neuroscience of Consciousness, 2016(1), niw001. https://doi.org/10.1093/nc/niw001

Rahnev, D. A., Bahdo, L., Lange, F. P. de, & Lau, H. (2012). Prestimulus hemodynamic activity in dorsal attention network is negatively associated with decision confidence in visual perception. Journal of Neurophysiology, 108(5), 1529–1536. https://doi.org/10.1152/jn.00184.2012

Rahnev, D., Maniscalco, B., Graves, T., Huang, E., de Lange, F. P., & Lau, H. (2011). Attention induces conservative subjective biases in visual perception. Nature Neuroscience, 14(12), 1513–1515. https://doi.org/10.1038/nn.2948

Rajagovindan, R., & Ding, M. (2010). From Prestimulus Alpha Oscillation to Visual-evoked Response: An Inverted-U Function and Its Attentional Modulation. Journal of Cognitive Neuroscience, 23(6), 1379–1394. https://doi.org/10.1162/jocn.2010.21478

Romei, V., Brodbeck, V., Michel, C., Amedi, A., Pascual-Leone, A., & Thut, G. (2008). Spontaneous Fluctuations in Posterior α-Band EEG Activity Reflect Variability in Excitability of Human Visual Areas. Cerebral Cortex, 18(9), 2010–2018. https://doi.org/10.1093/cercor/bhm229

Romei, V., Gross, J., & Thut, G. (2010). On the Role of Prestimulus Alpha Rhythms over Occipito-Parietal Areas in Visual Input Regulation: Correlation or Causation? The Journal of Neuroscience, 30(25), 8692–8697. https://doi.org/10.1523/JNEUROSCI.0160-10.2010

Romei, V., Rihs, T., Brodbeck, V., & Thut, G. (2008). Resting electroencephalogram alpha-power over posterior sites indexes baseline visual cortex excitability. Neuroreport, 19(2), 203–208. https://doi.org/10.1097/WNR.0b013e3282f454c4

Samaha, J. (2015). How best to study the function of consciousness? Consciousness Research, 6, 604. https://doi.org/10.3389/fpsyg.2015.00604

Samaha, J., Barrett, J. J., Sheldon, A. D., LaRocque, J. J., & Postle, B. R. (2016). Dissociating Perceptual Confidence from Discrimination Accuracy Reveals No Influence of Metacognitive Awareness on Working Memory. Consciousness Research, 851. https://doi.org/10.3389/fpsyg.2016.00851

Samaha, J., Gosseries, O., & Postle, B. R. (2016). Distinct oscillatory frequencies underlie excitability of human occipital and parietal cortex. bioRxiv, 82693. https://doi.org/10.1101/082693

Samaha, J., Sprague, T. C., & Postle, B. R. (2016). Decoding and Reconstructing the Focus of Spatial Attention from the Topography of Alpha-band Oscillations. Journal of Cognitive Neuroscience, 28(8), 1090–1097. https://doi.org/10.1162/jocn_a_00955

Sanders, J. I., Hangya, B., & Kepecs, A. (2016). Signatures of a Statistical Computation in the Human Sense of Confidence. Neuron, 90(3), 499–506. https://doi.org/10.1016/j.neuron.2016.03.025

Sauseng, P., Klimesch, W., Stadler, W., Schabus, M., Doppelmayr, M., Hanslmayr, S., … Birbaumer, N. (2005). A shift of visual spatial attention is selectively associated with human EEG alpha activity. European Journal of Neuroscience, 22(11), 2917–2926. https://doi.org/10.1111/j.1460-9568.2005.04482.x

Taylor, M. M., & Creelman, C. D. (1967). PEST: Efficient Estimates on Probability Functions. The Journal of the Acoustical Society of America, 41(4A), 782–787. https://doi.org/10.1121/1.1910407

Zoefel, B., & Heil, P. (2013). Detection of Near-Threshold Sounds is Independent of EEG Phase in Common Frequency Bands. Frontiers in Psychology, 4. https://doi.org/10.3389/fpsyg.2013.00262

Zylberberg, A., Barttfeld, P., & Sigman, M. (2012). The construction of confidence in a perceptual decision. Frontiers in Integrative Neuroscience, 6. https://doi.org/10.3389/fnint.2012.00079

Zylberberg, A., Roelfsema, P. R., & Sigman, M. (2014). Variance misperception explains illusions of confidence in simple perceptual decisions. Consciousness and Cognition, 27, 246–253. https://doi.org/10.1016/j.concog.2014.05.012

